# Molecular Characterization of T-Lineage Acute Lymphoblastic Leukemia by an Optimal-Transport Based Multi-Omics Integration Framework

**DOI:** 10.64898/2026.05.22.727257

**Authors:** Lusheng Li, Jieqiong Wang, Shibiao Wan

**Affiliations:** Department of Genetics, Cell Biology and Anatomy, University of Nebraska Medical Center, Omaha, NE, United States; Department of Neurological Sciences, University of Nebraska Medical Center, Omaha, NE, United States; Fred and Pamela Buffett Cancer Center, University of Nebraska Medical Center, Omaha, NE, United States

## Abstract

T-lineage acute lymphoblastic leukemia (T-ALL) is an aggressive pediatric malignancy characterized by complex heterogeneity across multiple molecular layers. Accurate subtyping is essential for understanding disease mechanisms, risk stratification, and guiding targeted therapeutic strategies. However, current diagnostic approaches are labor-intensive and time-consuming, and existing computational methods are limited by reliance on single-modality data or simple integration strategies. Effective integration of heterogeneous multi-omics data remains a major computational challenge. We present **OTTER** (**O**ptimal **T**ransport-based **T**ranscriptomics and g**E**nomics **R**epresentation), a novel multi-modal deep learning framework that jointly models transcriptomic and genomic data for T-ALL molecular characterization. OTTER encodes each omics modality through a modality-specific encoder and aligns the resulting latent representations using Gromov-Wasserstein optimal transport (GW-OT), which preserves the internal geometric structure of each modality without requiring a shared feature space. Benchmarking tests on the Children’s Oncology Group (COG) AALL0434 cohort with >1,300 patients across 17 T-ALL subtypes suggested that OTTER significantly outperformed state-of-the-art approaches for T-ALL molecular characterization. In addition, OTTER could identify interpretable subtype-defining and omics-specific features for T-ALL. Furthermore, we have demonstrated the generalizability of OTTER by evaluating on multiple external independent clinical cohorts. In summary, OTTER provides a principled, biologically interpretable, and computationally effective framework for multi-omics driven T-ALL molecular characterization and broader cancer applications.

## Background

T-lineage acute lymphoblastic leukemia (T-ALL) is an aggressive pediatric malignancy driven by coordinated alterations across multiple molecular layers, including epigenomics, genomics and transcriptomics (1,2). It originates from the malignant transformation of T-cell progenitors driven by uncontrolled proliferation, aberrant activation of oncogenic transcriptional programs, dysregulated signaling pathways, and disrupted T-cell developmental processes (3–5). T-ALL can be classified into 17 molecular subtypes based on integrative analyses of whole-transcriptome and whole-genome sequencing (WTS/WGS) data (2). These subtypes are characterized by distinct oncogenic drivers and co-occurring genomic alterations, correspond to specific stages of T-cell development, and are associated with differences in clinical outcomes. Accurate molecular subtyping is essential for understanding disease mechanisms, risk stratification, and the development of targeted therapeutic strategies (6–8). However, conventional methods for T-ALL subtype identification, including immunophenotyping (9,10), cytogenetic analysis (11), fluorescence in situ hybridization (FISH) (12), and targeted molecular assays (13,14), are often labor-intensive, time-consuming, and costly, and may provide only a partial view of the underlying molecular heterogeneity. Moreover, many T-ALL cases harbor biologically relevant alterations in noncoding genomic regions, such as regulatory elements and epigenetic modifications (15,16), which are difficult to detect using standard diagnostic workflows that primarily focus on coding mutations or predefined targets. In addition, the technical complexity of these approaches presents significant barriers to their widespread implementation in clinical practice, particularly in resource-limited settings. The interpretation of molecular assays requires domain expertise and is often subject to inter-observer variability, reducing reproducibility and limiting scalability. These challenges underscore the need for computational frameworks that can systematically integrate diverse molecular signals while reducing analytical burden and improving robustness.

Recent advances in machine learning (ML) have enabled data-driven approaches to leukemia subtype prediction (17–21) and subtyping in other types of cancer (22–24), providing promising strategies to enhance our understanding of disease heterogeneity and to support improved risk stratification and patient outcomes. Specifically, recent studies have introduced RanBALL (18), a random projection-based model for B-ALL subtyping, and AttentionAML (19), an attention-based deep learning framework for AML classification. However, these existing computational methods for leukemia subtyping have largely focused on transcriptomic profiles, leveraging gene expression patterns to distinguish disease subtypes. While transcriptomics provides a powerful snapshot of cellular state and has enabled important advances in leukemia classification, reliance on a single molecular modality imposes inherent limitations. Gene expression reflects downstream consequences of regulatory processes but does not fully capture the upstream genetic and epigenetic mechanisms that drive leukemia initiation and progression (1,2,25,26).

Advances in next generation sequencing have enabled comprehensive multi-omics profiling (27–30), offering an opportunity to capture these noncoding and regulatory alterations through modalities such as genomics and transcriptomics. However, effectively integrating these heterogeneous data types remains a major computational challenge. A fundamental challenge arises from the heterogeneous nature of multiomics data (31–33). Different modalities exhibit unique statistical properties, dimensionalities, noise characteristics, and measurement scales. For example, genomic data are often sparse and region-based, whereas transcriptomic data are continuous and high-dimensional. These differences make direct comparison or naive fusion of modalities suboptimal and prone to bias, motivating the need for integration strategies that respect modality-specific characteristics. Beyond statistical heterogeneity, molecular processes are governed by complex, non-linear interactions across modalities, such as epigenetic regulation of gene expression or feedback between transcriptional and signaling pathways. Capturing these cross-modal dependencies requires models that go beyond linear correlations and explicitly account for structured, non-linear relationships. Existing multimodal approaches for leukemia subtyping relies on simple integration strategies. For example, TALLForest (34) integrated gene expression-based random forest model with genomic variants to derive consensus subtype predictions. Additionally, clinTALL (35) concatenated the latent representations from different omics for downstream tasks. As existing methods fail to fully exploit the complementary information across molecular layers, advanced computational frameworks capable of jointly modeling heterogeneous multi-omics data in T-ALL are critically needed.

To overcome these challenges, we present **OTTER** (**O**ptimal **T**ransport-based **T**ranscriptomics and g**E**nomics **R**epresentation), a novel multi-modal learning framework that integrates transcriptomics data and genomics data for accurate and cost-effective T-ALL subtyping. OTTER was developed using paired molecular profiles from 1,309 patients enrolled in the Children’s Oncology Group (COG) AALL0434 study. The framework encodes each omics modality through modality-specific encoders and aligns the resulting latent representations using Gromov-Wasserstein optimal transport (GW-OT) (36,37), which preserves the internal geometric structure of each modality without requiring a shared feature space or explicit cross-modal sample correspondences. Benchmarking against state-of-the-art single-modality and multi-modality integration methods demonstrated that OTTER achieves superior subtype classification performance. The ablation and sensitivity analyses supported the contributions of GW-OT alignment and variational regularization. Beyond prediction, OTTER provides a gradient-based interpretability module that quantifies feature importance for each omics modality, enabling the identification of transcriptomic and genomic drivers of T-ALL subtypes. We demonstrate that the fused multimodal representation achieves superior subtype discrimination compared to either modality alone, recovers biologically established oncogenic features as top predictors, and reveals cross-modal interaction patterns consistent with known T-ALL pathobiology. Application to two independent clinical cohorts provided evidence that the learned representation is transferable across datasets. The OTTER framework is generalizable beyond T-ALL and offers a principled computational strategy for multi-omics data integration in any cancer type characterized by heterogeneous molecular alterations.

## Methods

### Data sources

The dataset was derived from Children’s Oncology Group (COG) AALL0434 trial (2) and obtained through the Synapse platform (ID: syn54032669; https://doi.org/10.7303/syn54032669). It comprises 1,309 patients that were identified as 17 T-ALL subtypes. This study integrated gene expression and DNA sequence variants data. **Fig. 1A** summarizes the clinical and biological characteristics of the 1,309 patients included in the COG AALL0434 cohort. Each column represents an individual patient, with patients grouped according to the 17 molecular T-ALL subtypes. The annotation tracks indicate molecular subtype, ETP phenotype, age at diagnosis, sex, central nervous system (CNS) status, and day-29 minimal residual disease (MRD). The cohort displayed a markedly imbalanced subtype distribution, with TAL1 DP-like (n = 296), ETP-like (n = 235), TAL1 αβ-like (n = 219), and TLX3 (n = 212) representing the largest groups. In contrast, several subtypes are comparatively uncommon, including NUP214 (n = 5), NUP98 (n = 6), and NKX2-5 (n = 8). ETP and near-ETP phenotypes were enriched among molecularly immature groups but were not completely restricted to a single genomic subtype, illustrating the distinction between immunophenotypic and molecular classifications. Patients showed a broad age distribution and an overall predominance of male patients across most subtypes. Most samples were classified as CNS1 at diagnosis, with smaller proportions assigned to CNS2 or CNS3. Day-29 MRD levels were heterogeneous both within and across molecular subtypes, ranging from undetectable disease to MRD levels of at least 1%.

**Figure 1.**
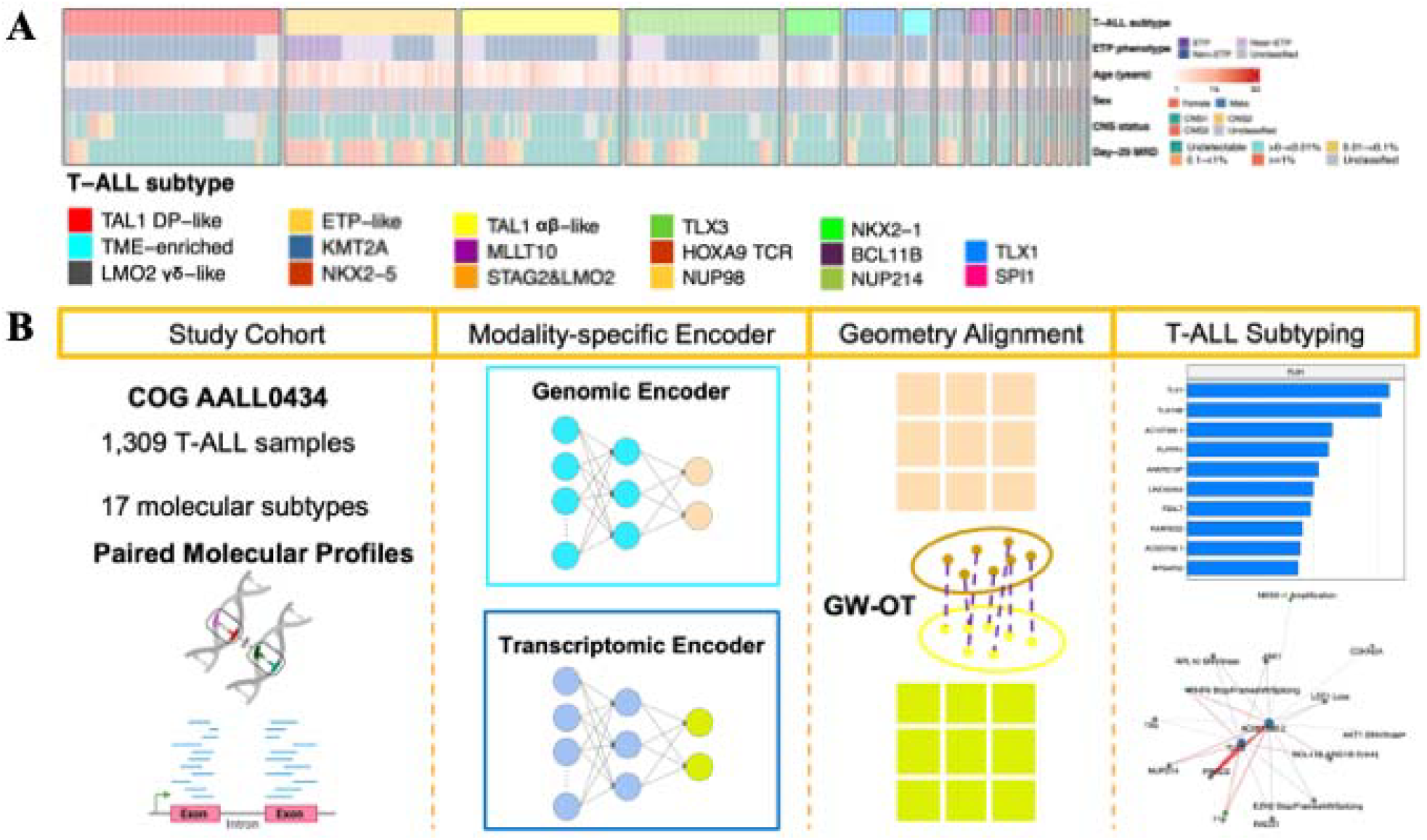
The overview of the proposed OTTER for T-ALL subtyping. **(A)** Clinical landscape of the T-ALL cohort. **(B)** Schematic overview of the proposed OTTER framework. Genomic and transcriptomic data are first processed through modality-specific encoders to extract latent representations for each modality. These modality-specific latent spaces are then aligned using GW-OT, which preserves relational structure across modalities. The aligned representations are subsequently integrated to generate a unified embedding for accurate T-ALL subtype classification.

### Data Preprocessing

Samples are first intersected across the two modalities to retain only the common set. Features are then scaled: Min-Max normalization is applied for classification tasks. Scaling parameters are estimated exclusively on the training partition and applied to both training and holdout sets to prevent data leakage. The dataset is split into training and holdout (test) partitions with stratified random split (80%/20%). The stratification ensures that class proportions are preserved in both partitions.

### OTTER framework

The novel Optimal Transport based multimodal representation learning framework that jointly models genomic and transcriptomic data was shown in **Fig. 1C**. The framework takes genomics and transcriptomics data modalities as input. Due to the fundamentally different statistical properties of these modalities, each is first processed by a modality-specific encoder designed to extract biologically meaningful features. Each omics modality (m ∈ A,B) is processed by a two-layer feed-forward encoder:

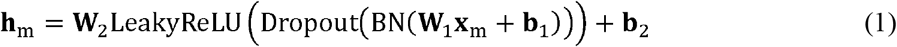

where batch normalization (BN) and dropout are applied after the first linear layer. LeakyReLU activation functions (38) were applied after the hidden linear layers to introduce nonlinearity while retaining a small, nonzero gradient for negative input values. The encoded features are then projected into latent spaces (default 100) using modality-specific encoder, enabling the learning of compact, informative embeddings for each modality. The encoder output is projected to Gaussian parameters via two parallel linear heads:

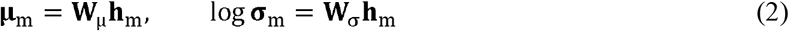

During training, a latent sample is drawn by the reparameterization trick:

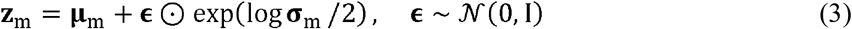

During inference, **z**_m_ = μ_m_.

Each modality’s latent distribution is regularized toward (*N*(0,*I*)):

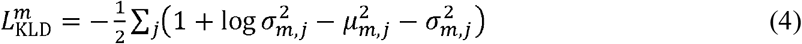

The core component of framework is an optimal transport-based alignment module. Optimal Transport seeks an optimal correspondence between two distributions by minimizing the cost of transporting probability mass while preserving their underlying structure (39). Specifically, we employ Gromov-Wasserstein optimal transport (GW-OT) (40) to align the latent representations of the genomic and transcriptomic embeddings extracted from encoder. Unlike conventional alignment strategies that rely on direct feature or sample-level correspondence, GW-OT operates on pairwise relational structures, aligning the two modalities by preserving their internal geometric relationships. The OT loss penalizes the discrepancy between these two latent distributions and is incorporated into the total training objective alongside the task loss and KL divergence terms. This enables the identification of shared structural patterns across modalities, even when their feature spaces differ substantially. Following GW-based alignment, the model produces unified latent representations that capture complementary information from both genomics and transcriptomics. These aligned embeddings are then integrated and provided as input to a downstream classifier, which performs T-ALL subtype prediction.

### Optimal Transport Alignment

To align the latent representations learned from the two omics modalities, we adopt GW-OT as the cross-modal alignment objective. Unlike conventional OT formulations that require a shared ambient feature space and direct cross-modal distances, GW-OT operates on the internal geometry of each latent space independently, making it particularly well-suited for heterogeneous omics data where the two modalities encoders produce latent Gaussian parameters (**μ**_A_, log **σ**_A_) and (**μ**_B_, log **σ**_B_)for omics A and B, may reside in spaces of different nature and dimensionality. After each forward pass, the variational respectively. Rather than penalizing direct distances between 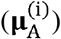 and 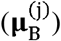, which would implicitly assume the two latent spaces are directly comparable, GW-OT instead seeks a transport plan that preserves relational structure: pairs of samples that are close in the omics A latent space should be mapped to pairs of samples that are close in the omics B latent space.

For each modality, we compute the pairwise Euclidean distance matrix over the latent means within the mini batch:

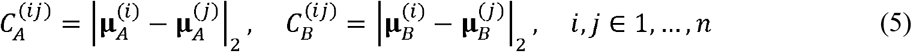

Both matrices are then normalized by their respective maxima to bring them to a common scale:

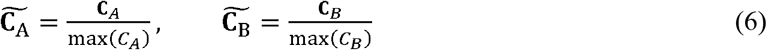

This normalization ensures that the GW alignment is not dominated by differences in the absolute scale of distances between the two modalities.

Given uniform marginal distributions (p = 1_n_/n) over omics A samples and (q **1**_n_/n) over omics B samples, the GW transport plan Γ_GW_ ∈ ℝ^nxn^ is obtained by solving the following quadratic program:

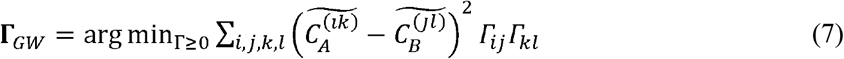

subject to the marginal constraints:

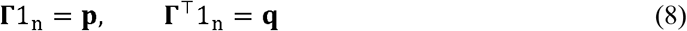

This objective minimizes the structural discrepancy between the two latent spaces: a small GW cost means that the relational geometry of omics A samples is faithfully reflected in the relational geometry of their matched counterparts in omics B. The problem is solved under the square-loss formulation using the POT (Python Optimal Transport) library.

The GW alignment loss incorporated into the total training objective is:

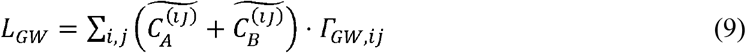

The transport plan (**Γ**_GW_) is detached from the computational graph (treated as a fixed coupling matrix), so gradients during backpropagation flow exclusively through the intra-modality distance matrices 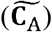 and 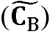, and consequently through the encoder parameters (**μ**_A_) and (**μ**_B_). This encourages the encoders to organize their latent spaces so that the relational structure between samples is consistent across the two omics.

The GW loss is combined with the variational KL divergence terms and the task-specific classification via an adaptive weighting scheme:

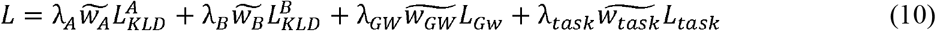

where 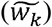 are dynamic inverse-magnitude normalization factors that prevent any single loss component from dominating training. The GW-OT term thus acts as a geometry-preserving regularizer, aligning the two latent spaces in a manner that is agnostic to the original feature dimensionalities of the input omics.

### Cross-Validation and Training Procedure

Model performance is evaluated using K-fold cross-validation on the training partition (default (K =5)). Stratified K-Fold is used to preserve class proportions across folds. Within each fold, a validation subset is held out for monitoring. Model parameters are optimized using the Adam optimizer. A learning rate scheduler and early stopping (monitored on validation classification loss, default patience = 40 epochs) are applied to prevent overfitting. The model state that achieves the lowest validation loss is retained for evaluation on the holdout set at the end of each fold. Reported metrics include Accuracy, Precision, Recall, F1 score, and Matthews Correlation Coefficient (MCC).

### Gradient-based Feature Importance Analysis

Gradient-based feature attribution (41) was used to quantify the contribution of individual transcriptomic and genomic features to subtype classification. After training, gradient-based feature importance scores are computed on the holdout set.

For each holdout sample, the model output is backpropagated to the input features of each omics. The objective is the sum of the true-class logit across samples. The per-feature importance for omics (m) is:

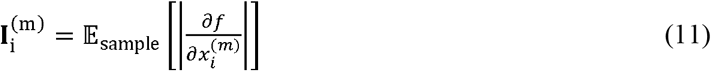

where 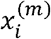is the value of feature i in modality m for a given sample after preprocessing. Gradients are taken with respect to these scaled inputs. Scalar objective *f* derived from the trained OTTER model output on holdout samples. It is the quantity whose sensitivity to inputs we measure sum of the logit for the true class label for each sample.

### Subtype-Specific Differential gene Expression Analysis

Subtype-specific differential gene-expression analysis was performed using the limma R package v3.54.2 (42). The analysis used the normalized, log-transformed expression matrix from the 1,309 samples with molecular subtype annotations. For each of the 17 T-ALL subtypes, a one-versus-rest contrast compared samples belonging to the target subtype with all remaining samples. Moderated test statistics were estimated using empirical Bayes moderation with mean-variance trend estimation. Multiple-testing correction was performed separately for each subtype using the Benjamini-Hochberg method. Genes with an adjusted P value below 0.05 and a log2 fold change of at least 1 were classified as significantly upregulated, whereas genes with an adjusted P value below 0.05 and a log2 fold change of lower −1 were classified as significantly downregulated.

### Subtype-Specific Genomic-Feature Analysis

For each subtype and genomic feature, a 2 × 2 table summarized the presence or absence of the feature in samples belonging to the target subtype and in all remaining T-ALL samples. Positive subtype enrichment was evaluated using a one-sided Fisher’s exact test with the alternative hypothesis that the genomic event was more prevalent in the target subtype. Feature prevalence within the subtype, prevalence in the remaining cohort, and the corresponding prevalence difference were also calculated. To obtain finite effect-size estimates for tables containing zero cells, odds ratios were calculated using the Haldane-Anscombe correction, in which 0.5 was added to each cell, and were subsequently expressed as log2 odds ratios. Fisher-test P values were adjusted within each subtype using the Benjamini-Hochberg method. Features with an adjusted P value below 0.05 and a positive log2 odds ratio were considered significantly enriched.

## Results

### OTTER Outperforms Other State-of-the-art (SOTA) Methods in T-ALL subtype prediction

We developed **OTTER** (Optimal Transport-based Transcriptomics and gEnomics Representation), an accurate and cost-effective model to identify T-ALL subtypes based on genomics and transcriptomics data (**Fig. 2**). OTTER was designed to address the limitations of conventional diagnostic approaches and single-omics computational models by integrating transcriptomic and genomic data through a classical Optimal Transport alignment strategy (43). In this framework, genomic and transcriptomic profiles were first processed through encoder to extract informative modality-specific features. Optimal Transport was then employed to align and fuse heterogeneous omics representations by learning a cross-modal cost matrix that quantifies alignment between modalities, thereby capturing complementary biological interdependencies. To evaluate the performance of OTTER, 20% hold-out test was conducted for the T-ALL dataset. The **Fig. 2A-B** summarizes the comparative performance of multiple computational methods for T-ALL subtype prediction across five standard evaluation metrics: Accuracy, Precision, Recall, F1-score, and MCC. The baseline methods include single-modality approaches (AttenTALL (19), RanTALL (18), and TALLSorts (17)) and multi-modality approach clinTALL (35). As shown in **Fig. 2A**, OTTER consistently outperforms existing single-modality approach AttenTALL with RNA and SNV data across all evaluation metrics. OTTER achieves the highest median performance with lower variability, indicating improved robustness and generalization. For the multi-modality approach (**Fig. 2B**), OTTER demonstrates consistently higher performance across all metrics compared to the clinTALL. Notably, the strongest improvement is observed in MCC, a balanced metric that accounts for class imbalance and false predictions, underscoring OTTER’s robustness in leukemia subtyping settings. These results provide preliminary evidence that explicit cross-omics alignment using Optimal Transport yields a more informative and stable latent representation than heuristic or single-modality approaches. By jointly modeling shared and complementary biological signals across data modalities, OTTER improves predictive performance and consistency across evaluation criteria.

**Figure 2.**
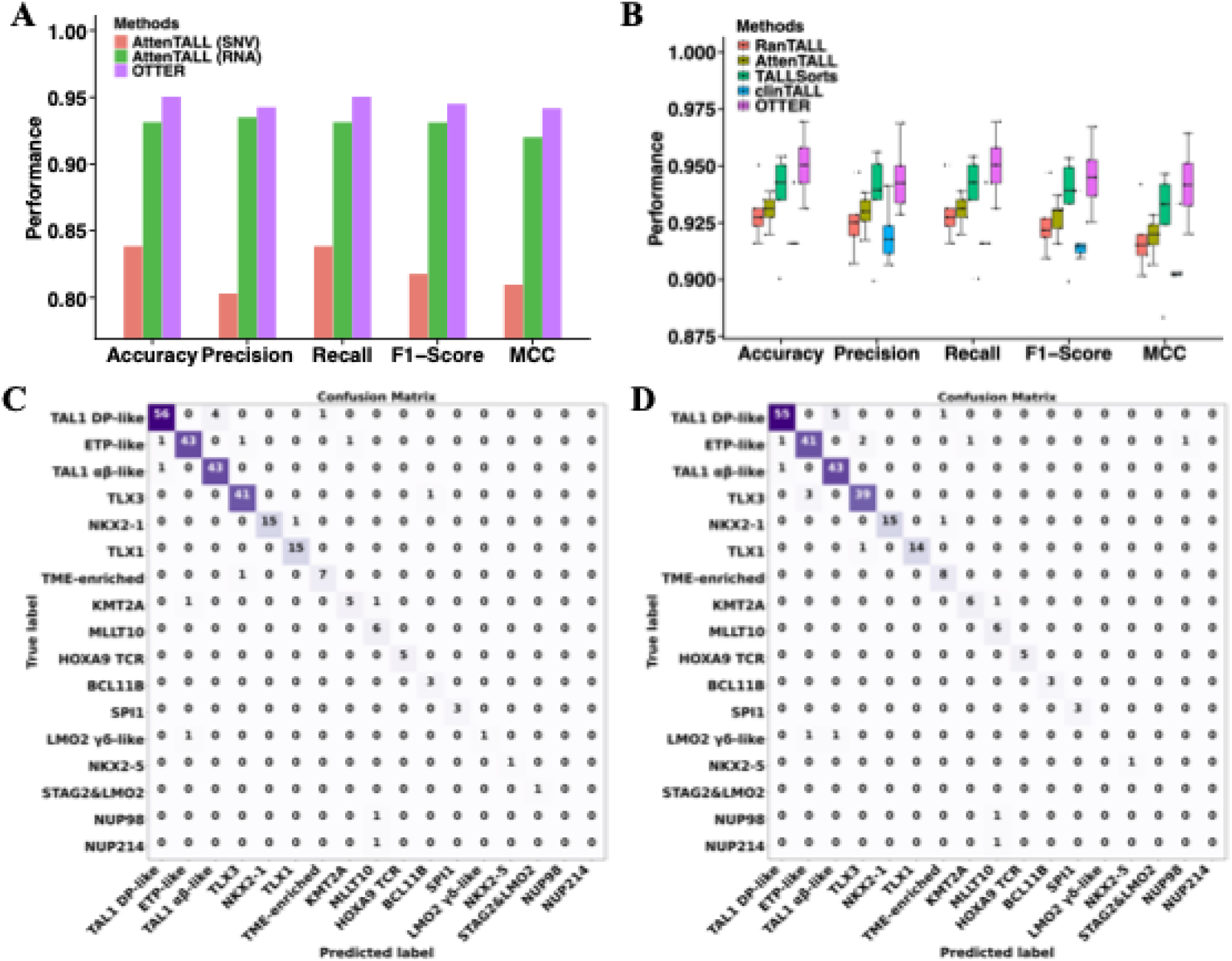
Performance comparison on T-ALL classification. **(A)** OTTER outperformed state-of-the-art methods like AttenTALL. **(B)** Integrating multi-omics data by OTTER performed better than using SNV data or RNA-seq data alone for T-ALL subtyping. **(C)** Confusion matrix of OTTER on the 20% hold-out test set. **(D)** Confusion matrix of clinTALL model on the same hold-out test set.

To further validate the OTTER model, we assessed its performance on an independent 20% hold-out test set derived from the T-ALL dataset. To further investigate prediction confidence, **Fig. 2B** visualizes the distribution of predicted probabilities across T-ALL subtypes. OTTER produces highly confident predictions for the majority of samples, with probabilities concentrated near 1.0, while misclassified or uncertain samples exhibit low probabilities. Notably, confidence varies across subtypes, with rarer or more heterogeneous groups (e.g., STAG2&LMO2, NUP98, NUP214) showing increased dispersion, reflecting intrinsic classification difficulty. The confusion matrices in **Fig. 2C-D** provide a detailed view of subtype-specific performance. OTTER (**Fig. 2C**) demonstrates strong diagonal dominance, indicating accurate classification across most subtypes, particularly for major groups such as TAL1 DP-like, ETP-like, and TAL1 αβ-like. In comparison, the clinTALL model (**Fig. 2D**) exhibits more frequent off-diagonal errors, especially for closely related or underrepresented and phenotypically similar subtypes, indicating weaker generalization and reduced discriminative power.

### Identification of subtype-associated transcriptomic and genomic drivers via gradient-based feature importance

Gradient-based feature importance analysis revealed distinct subtype-associated molecular patterns across both transcriptomic and genomic modalities (**Fig. 3**). The subtype-specific heatmaps demonstrated that many of these features formed discrete attribution programs, while a subset was shared across related subtypes, suggesting that the model integrated both subtype-defining drivers and broader developmental or signaling programs.

**Figure 3.**
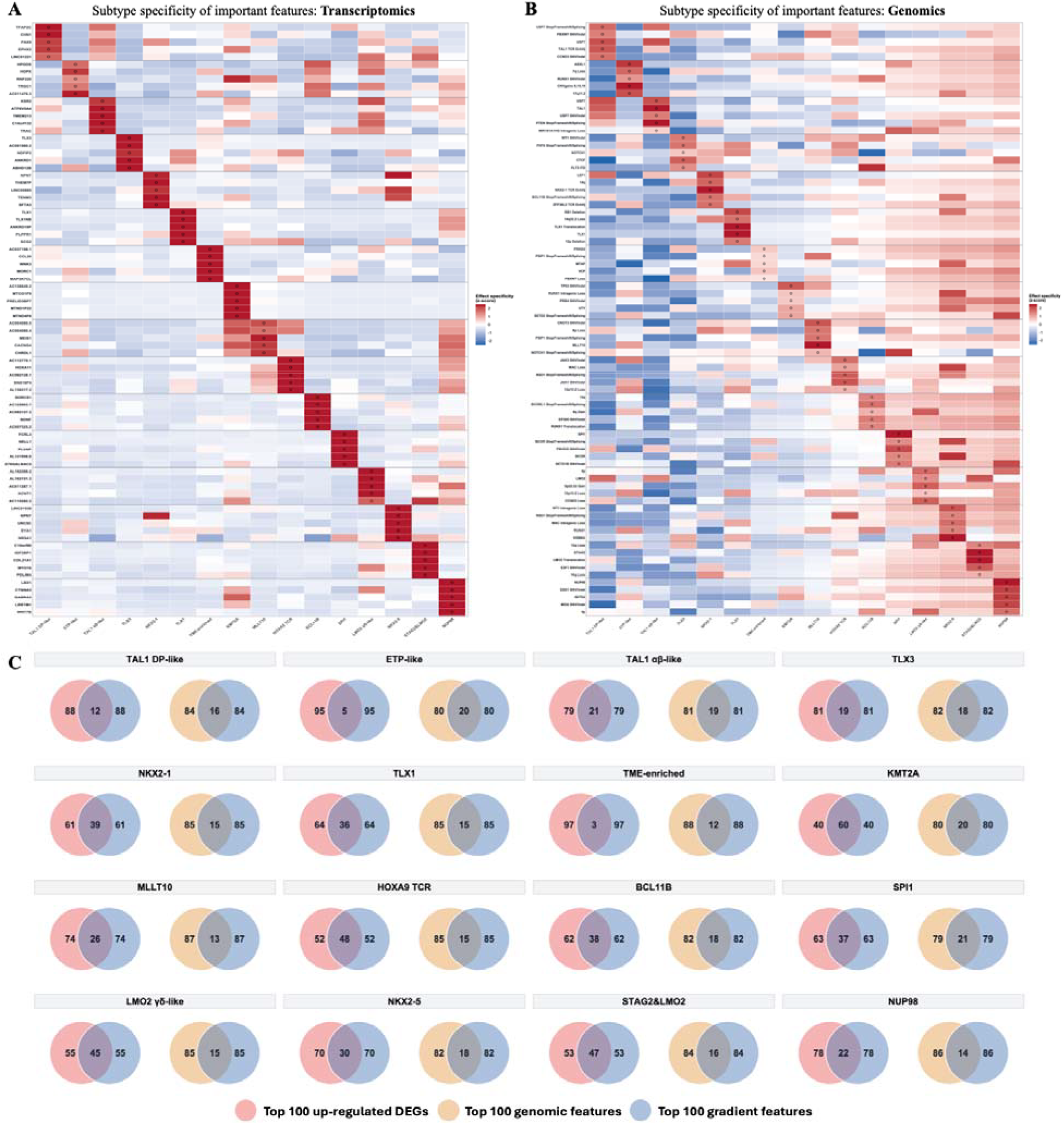
Gradient-based feature importance landscape for T-ALL subtype classification. **(A)** Subtype specificity of transcriptomic features shared between the top-ranked gradient features and upregulated differentially expressed genes. **(B)** Subtype specificity of genomic features shared between conventional genomic feature selection and gradient-based attribution. Rows represent selected transcriptomic features and columns represent T-ALL molecular subtypes. Cell colors indicate standardized effect-specificity scores, with red denoting higher positive subtype specificity and blue denoting lower or negative specificity. Black circles identify features ranked among the top 5 within the corresponding subtype. (**C**) Venn diagrams comparing conventional and gradient-based feature rankings for each subtype. For every subtype, the left diagram compares the top 100 upregulated differentially expressed genes with the top 100 transcriptomic gradient features, whereas the right diagram compares the top 100 conventionally selected genomic features with the top 100 genomic gradient features.

In the transcriptomic modality, the model recovered several biologically recognizable subtype-defining expression programs (**Fig. 3A**). TLX3 and TLX1/TLX1NB were among the most important features for the corresponding TLX3 and TLX1 subtypes. This is consistent with established biology: TLX3 is canonical oncogenic driver that define distinct T-ALL subtypes (44–47). Similarly, NKX2-1 and its antisense transcript NKX2-1-AS1 were the two most influential transcriptomic features for NKX2-1classification. These findings indicate that the model relied directly on the expression of the defining transcription factor and associated transcriptional programs for these subtypes. This observation is consistent with the established role of rearrangement-driven activation of TLX1, TLX3, and NKX2-family transcription factors in defining molecular subtypes of T-ALL. In particular, integrated genomic and transcriptomic analyses previously identified rearrangement-associated NKX2-1 activation as a defining event in a distinct T-ALL subtype (48). The MLLT10 subtype showed a recognizable HOXA/MEIS-associated transcriptomic pattern. The prominence of multiple MEIS1-associated antisense transcripts is consistent with the activation of HOXA/MEIS transcriptional programs commonly observed in leukemias driven by MLLT10 and related HOXA-activating rearrangements. Other T-ALL subtypes exhibited less canonical but clearly distinguishable patterns. SPI1 (**Fig. 3G**) predictions were associated with FCRL4, ST6GALNAC5 and PLVAP, consistent with the possibility that the classifier recognized the broader nonconventional hematopoietic or progenitor-like state associated with SPI1-rearranged leukemia rather than SPI1 expression alone. SPI1 rearrangements have been reported to define a distinct pediatric T-ALL subgroup with an immature phenotype and adverse outcome (49). These features should be considered as candidate subtype-associated expression signatures.

In the genomic modality, subtype predictions were informed by distinct combinations of structural variants, enhancer alterations, sequence variants, copy-number changes, and chromosomal gains or losses (**Fig. 3B**). For example, the diversity of these features for TLX3 subtype indicates that the model combined a subtype-defining transcriptional state with alterations affecting differentiation, epigenetic regulation, RAS/kinase signaling, and NOTCH signaling. In NKX2-1 and TLX1, the transcriptomic modality provided direct evidence of TLX or NKX2 oncogene activation, whereas the genomic modality predominantly captured accompanying chromosomal and cooperating alterations. Within the HOXA-associated groups, although not every high-ranking feature is itself the subtype-defining driver, the recurrence of enhancer-hijacking events and alterations involving HOXA-associated regulatory programs supports the model’s use of biologically related genomic contexts. Large-scale studies have shown that noncoding and enhancer-rewiring events are major mechanisms of oncogene deregulation in T-ALL (2). Overall, these results demonstrate that the model not only achieves high predictive performance but also captures biologically meaningful multimodal interactions, providing biologically interpretable insights into the molecular mechanisms underlying T-ALL subtypes. It combined direct subtype-defining signals with downstream expression programs and cooperating genomic abnormalities. This pattern is consistent with the molecular complexity of T-ALL, in which transcription-factor activation, developmental arrest, enhancer deregulation, signaling abnormalities, and tumor-suppressor loss jointly define T-ALL subtype.

To assess agreement between model-derived feature importance and conventional molecular analyses, we compared the top 100 gradient-ranked features with the top 100 upregulated differentially expressed genes and independently selected genomic features for each subtype (**Fig. 3C**). Transcriptomic overlap varied substantially among subtypes, ranging from 3 to 60 genes. The greatest agreement was observed for KMT2A, with 60 overlapping genes, followed by HOXA9 TCR with 48, STAG2&LMO2 with 47, LMO2 γδ-like with 45, and NKX2-1 with 39. For the genomics, the largest genomic overlaps were observed for SPI1, ETP-like, and KMT2A, with 21, 20, and 20 shared features, respectively. The overlapping features provide support for the biological relevance of the model explanations, whereas the non-overlapping gradient-ranked features suggest that OTTER also captures cross-omics predictive signals that may be missed by conventional approaches.

### Latent Space Visualization Demonstrates Improved Subtype Discrimination through OTTER

To visualizes the latent representations learned by the OTTER framework for two different data modalities, the PCA (50) and t-SNE (51) for dimensionality reduction was leveraged (**Fig. 4**). Each point represents a T-ALL sample, colored by its molecular subtype. The PCA was used to assess the global, linearly captured structure of the modality-specific and fused latent representations (**Fig. 4A-C**). In the transcriptomic latent space (**Fig. 4A**), several major T-ALL subtypes, icluding TAL1 DP-like, TAL1 αβ-like, ETP-like, TLX1, and TLX3, occupied distinguishable regions. In contrast, the genomic latent representation showed extensive mixing and little subtype-specific organization in the first two principal components (**Fig. 4B**). Following multi-omics integration, the fused representation retained the strong subtype-associated structure contributed by transcriptomic data while improving the compactness and separation of several subtype groups (**Fig. 4C**). In particular, TAL1 DP-like, TAL1 αβ-like, ETP-like, TLX1, and TLX3 formed more coherent regions, whereas samples belonging to less abundant subtypes remained primarily distributed between the major clusters. These patterns also were shown in the t-SNE visualization (**Fig. 4D-F**). In the transcriptomics latent space (**Fig. 4D**), samples form well-separated and compact clusters corresponding to T-ALL subtypes, such as TAL1 DP-like, ETP-like, TLX1, TLX3, and HOXA9 TCR. The clear separation between clusters indicates that the transcriptomic encoder learns a highly discriminative latent representation that captures subtype-specific gene expression programs. In contrast, the genomics latent space exhibits a more diffuse and overlapping structure (**Fig. 4E**). This reflects the inherent sparsity and heterogeneity of mutation data, where individual variants alone provide limited discriminatory power across subtypes. While transcriptomic data provide strong subtype discrimination, genomic variant data alone are less separable, motivating the need for cross-omics integration. Compared with the modality-specific latent spaces, the fused representation exhibits markedly improved cluster separation and compactness across subtypes (**Fig. 4F**). Distinct T-ALL subtypes form well-defined, non-overlapping clusters, indicating that the fusion process effectively integrates complementary information from different modalities while suppressing modality-specific noise. Samples from the same subtype form compact and well-isolated clusters, while inter-subtype overlap is substantially reduced. For example, TAL1 αβ-like shows markedly improved separation from neighboring TAL1-associated subtypes, suggesting that OTTER better captures subtle subtype-specific molecular differences within the TAL1 lineage. In addition, ETP-like appears more localized and separated in the OTTER visualization compared with the fused representation. Subtypes with limited number of patients, such as TME-enriched and KMT2A, forms more distinct and compact cluster with clear spatial separation from other subtypes compared with other representations. These results provide qualitative evidence that aligning and integrating heterogeneous latent spaces rather than relying on a single modality, can yield a more informative and biologically coherent representation for robust T-ALL subtype prediction.

**Figure 4.**
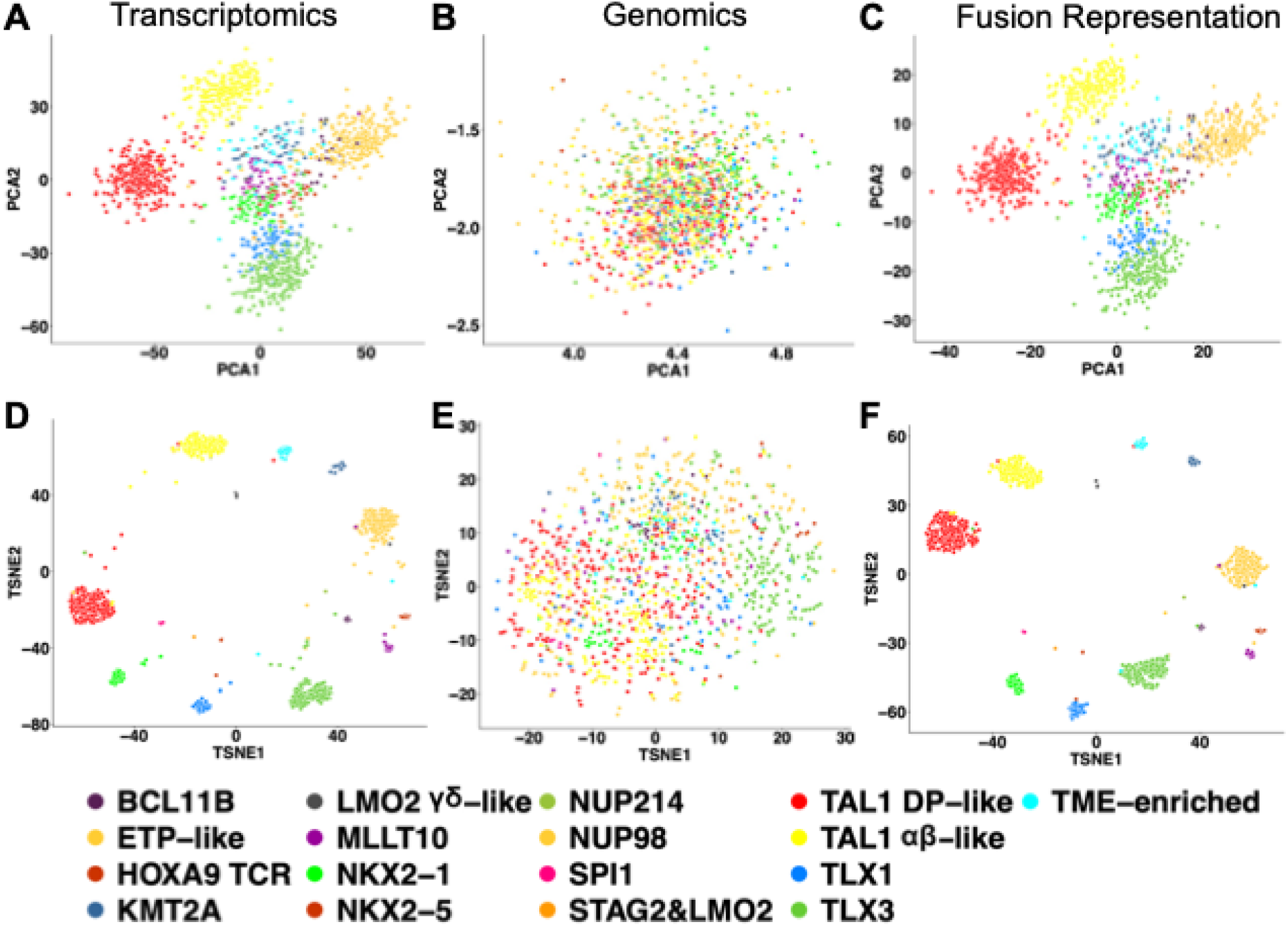
The visualization of latent spaces from OTTER framework. The PCA and t-SNE visualizations of latent embeddings learned from **(A, D)** transcriptomic data, **(B, E)** genomic data, and **(C, F)** the fused multimodal representation produced by the proposed OTTER framework. Each point represents a T-ALL sample and is colored by its molecular subtype.

### Ablation Studies Demonstrates the Effectiveness of OTTER

We first compared OTTER against three alternative strategies for aligning latent representations between the two omics views (**Fig. 5A**). Diagonal OT (52) enforces an index-aligned coupling that penalizes only paired within-batch distances, without estimating a transport plan. EMD (Earth Mover’s Distance) (53) computes a classical Wasserstein-1 alignment from pairwise latent costs, and entropic regularized OT (54) uses a Sinkhorn approximation for computational efficiency. OTTER consistently outperformed all three alternative strategies across all metrics, suggesting that both an explicit transport plan and a geometry-aware alignment in OTTER contribute to classification performance. We next quantified the contribution of optimal-transport alignment and VAE-style KL regularization by selectively removing these terms during training (**Fig. 5B**). Removing OT (w/o OT) decreased performance relative to the full model, demonstrating that cross-omics distributional alignment is beneficial beyond simple fusion of latent features. Removing KL regularization (w/o KLD) produced a larger drop, indicating that constraining latent representations remains important for stable learning and generalization. The largest degradation occurred when both OT and KL were disabled (w/o OT&KLD). The complete OTTER model recovered the highest performance on every metric, supporting the joint use of latent regularization and optimal-transport-based cross-omics coupling in the proposed framework.

**Figure 5.**
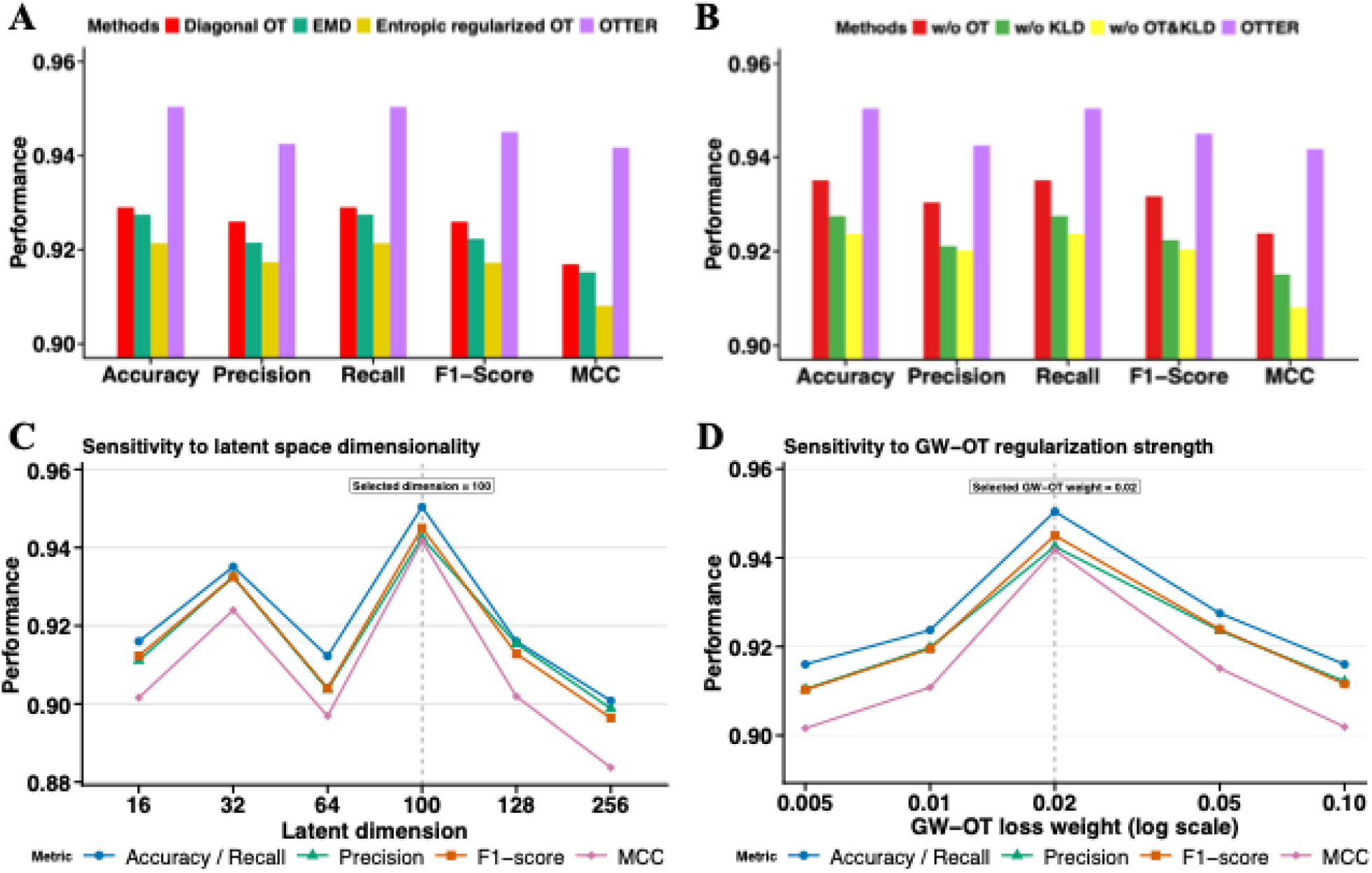
Ablation analysis of OTTER framework. **(A)** OTTER outperformed the alternative optimal-transport formulations for cross-omics alignment: Diagonal OT (index-aligned coupling), Earth Mover’s Distance (EMD), and entropic regularized OT (Sinkhorn). **(B)** Ablation of core regularization components in the full OTTER model: training without OT alignment (w/o OT), without variational KL penalties on either omics encoder (w/o KLD), without both terms (w/o OT&KLD), and the complete model (OTTER). (**C**) Sensitivity of OTTER to the dimensionality of the latent representations by dimensions. (**D**) Sensitivity of OTTER to strength of GW-OT regularization. Performance is evaluated by Accuracy, Precision, Recall, F1-score, and MCC.

In addition, we evaluated the sensitivity of OTTER to the dimensionality of the modality-specific latent representations by testing dimensions ranging from 16 to 256 (**Fig. 5C**). A latent dimension of 100 achieved the strongest performance across all evaluation metrics. A smaller latent dimension of 32 also performed comparatively well, whereas both very low and excessively high dimensionalities resulted in reduced performance. These results suggest that a moderately sized latent representation provides sufficient capacity to capture cross-omics information while limiting redundant dimensions and potential overfitting. Accordingly, a latent dimension of 100 was selected for the final OTTER model. For the strength of GW-OT regularization (**Fig. 5D**), increasing the GW-OT loss weight from 0.005 to 0.02 progressively improved all evaluated metrics, with a weight of 0.02 achieving the highest performance. Further increasing the weight to 0.05 or 0.10 reduced performance, indicating that excessive geometry alignment may constrain subtype-discriminative information specific to each modality. These results support a GW-OT weight of 0.02 as an appropriate balance between cross-modal alignment and preservation of modality-specific signals.

### OTTER Demonstrates Generalizability Across Independent T-ALL Cohorts

To evaluate whether the latent representation learned by OTTER generalized to independent clinical cohorts, we projected the training samples together with two external RNA-sequencing cohorts into a shared t-SNE space (**Fig. 6**). The training cohort (**Fig. 6A**) formed several well-defined subtype-associated regions, with particularly clear separation of the major TAL1 DP-like, TAL1 αβ-like, NKX2-1, TLX1, TLX3, ETP-like, KMT2A, MLLT10, and TME-enriched groups, whereas several rare subtypes occupied more closely positioned regions. The samples from NOPHO ALL2008 cohort (**Fig. 6B**) (55) assigned to the major TAL1 DP-like, TAL1 αβ-like, NKX2-1, TLX1, TLX3, and TME-enriched subtypes generally overlapped with or localized adjacent to the corresponding training regions, indicating preservation of subtype-associated latent structure across cohorts. The AIEOP cohort (**Fig. 6C**) (35) displayed a similarly subtype-dependent pattern: samples predicted as KMT2A and MLLT10 mapped closely to their corresponding reference clusters, while ETP-like and TAL1 αβ-like samples occupied the same upper region of the embedding observed in the training data. Predicted TAL1 DP-like, TME-enriched, TLX3, BCL11B, and several rarer subtype samples were positioned within or near the corresponding portions of the reference space. Compared with NOPHO ALL2008 cohort, however, the AIEOP samples were more concentrated in the central region. This compression may reflect differences in sequencing protocols, normalization, cohort composition, tumor purity, or batch effects, but the preservation of the overall relative subtype arrangement indicates that OTTER captured biological structure across independent datasets. Overall, the concordant positioning of predicted subtypes in both external cohorts (**Fig. 6D**) supports the transportability of the learned latent representation to real-world diagnostic RNA-sequencing data. Nevertheless, because ground-truth subtype annotations were unavailable for most external samples, these findings demonstrate consistency with the learned reference molecular representation.

**Figure 6.**
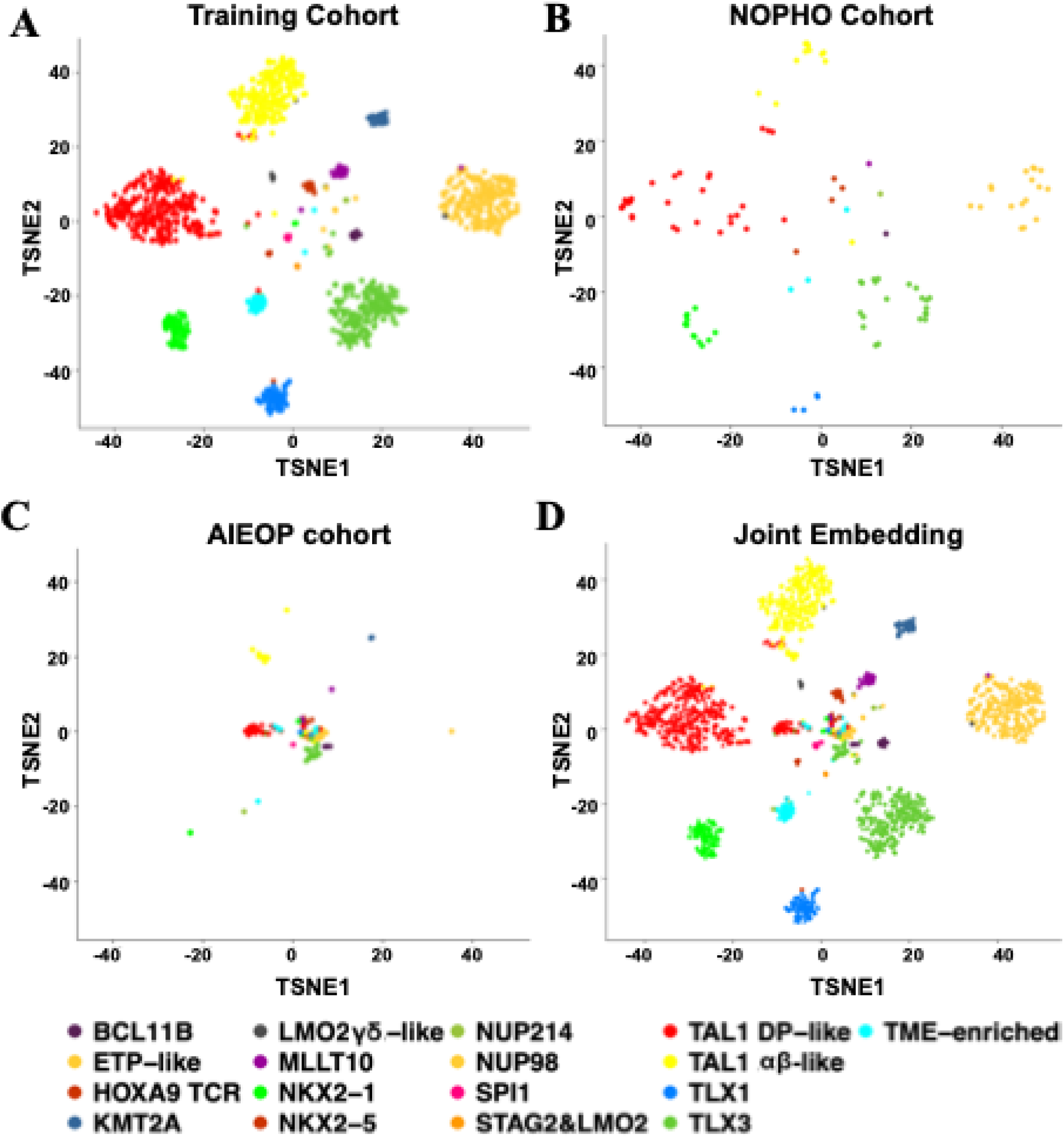
Evaluating and validating the generalizability of OTTER on multiple external independent T-ALL cohorts. The t-SNE visualization of OTTER latent representations, with each point representing one patient and colors denoting predicted T-ALL subtypes. **(A)** Training cohort. **(B)** Independent NOPHO ALL2008 cohort. **(C)** Independent AIEOP-BFM ALL 2017 cohort. **(D)** Joint visualization of the training and external cohorts in a shared latent space.

## Discussion

In this study, we presented OTTER, a multi-modal deep learning framework that integrates transcriptomics and genomic data for accurate molecular subtyping of T-ALL. By coupling modality-specific encoders with GW-OT alignment and an adaptive multi-component loss function, OTTER addresses several fundamental limitations of existing computational methods for leukemia classification. Recent computational approaches to T-ALL subtyping have largely relied on single-omics transcriptional classifiers, such as TALLSorts (17), which leverages RNA-seq expression profiles to assign subtypes. More recently, multi-modal approaches including clinTALL (35) and TALLForest (34) have begun to incorporate genomic information alongside transcriptomics. A central component in OTTER is the use of GW-OT to align the latent representations of the two omics modalities. Unlike conventional integration strategies, such as feature concatenation or correlation-based fusion, GW-OT does not require a shared ambient space or direct cross-modal feature correspondences. Instead, it aligns the two latent spaces by preserving their internal relational geometry, making it particularly well-suited for heterogeneous omics data where transcriptomic and genomic features differ fundamentally in their statistical properties, dimensionalities, and measurement scales.

OTTER consistently outperformed the evaluated single-modality and multimodal approaches (**Fig. 2**), supporting the premise that explicit integration of transcriptomic and genomic information improves T-ALL subtype classification. The strong improvement in MCC is important because this metric is less influenced by the imbalance among the 17 T-ALL subtypes than overall accuracy. The benchmarking results indicate that geometry-aware multi-omics integration provides a more informative representation than either modality alone or simpler multimodal fusion strategies.

The gradient-based feature importance analysis suggests that OTTER’s predictive performance is grounded in biologically feature programs (**Fig. 3**). Their importance scores confirm that OTTER successfully learns to exploit the most biologically meaningful transcriptional programs. Similarly, among genomic features further aligns with established knowledge of genome instability and structural variants in T-ALL pathogenesis. Overlapping features between gradient-ranked features and conventional differential gene expression analysis or genomic rankings provide internal evidence that the OTTER uses established subtype-associated signals. The identified features should consequently be regarded as hypothesis candidates requiring confirmation through independent cohorts, regulatory network analysis, and functional perturbation experiments.

The improved cluster separation observed in the fused latent space (**Fig. 4C, F**) compared to either individual modality alone (**Fig. 4A-B, D-E**) provides direct visual evidence that GW-OT alignment successfully captures complementary inter-modal structure that neither modality encodes on its own. This pattern is consistent with the intended role of GW-OT: rather than forcing direct equality between heterogeneous features, the alignment objective encourages the genomic and transcriptomic encoders to preserve comparable relationships among patients. The resulting representation can therefore incorporate complementary genomic context without discarding the discriminative transcriptional structure. This is especially important for genomic variant data, which are inherently sparse and provide limited discriminatory power when used in isolation, as reflected by the more diffuse clustering pattern in the genomics-only latent space.

The ablation experiments (**Fig. 5A-B**) clarify that OTTER’s performance reflects the joint effects of variational regularization and geometry-aware alignment. The superiority of index-aligned coupling (Diagonal OT), Earth Mover’s Distance (EMD), and entropic regularized OT (Sinkhorn) suggests that preserving within-modality relational geometry is advantageous when integrating data types that do not share a directly comparable feature space. The reduction in performance after removal of the OT term supports the contribution of cross-modal alignment beyond simple latent fusion. In addition, the decreased performance with removal of KL regularization indicates that constraining the modality-specific latent distributions is important for stable and generalizable learning. The sensitivity analyses (**Fig. 5C-D**) demonstrate that model performance depends on balancing representational capacity and alignment strength. Very small embeddings may omit subtype-relevant information, whereas excessively large embeddings may retain redundant or modality-specific noise. Similarly, insufficient GW weighting provides limited cross-modal coordination, while excessive weighting may suppress informative differences between the modalities.

Projection of the NOPHO ALL2008 and AIEOP-BFM ALL 2017 cohorts into the learned representation (**Fig. 6**) provides evidence that OTTER captures subtype-associated feature structure that is transferable across independent clinical datasets. External samples assigned to major predicted subtypes generally localized within or adjacent to the corresponding regions of the training reference. This pattern in two cohorts supports the robustness of the learned representation to differences in cohort composition and data source. The preservation of the broad subtype-associated structure across both cohorts supports the potential generalizability of OTTER to heterogeneous real-world T-ALL datasets.

Despite the aforementioned significant contributions of OTTER, we admit that this study might have some limitations. First, the COG AALL0434 cohort, while large, is highly imbalanced across the 17 T-ALL subtypes, with rare subtypes such as NUP214 (n=5), NUP98 (n=6), and NKX2-5 (n=8) comprising very few samples. This imbalance inevitably limits the model’s ability to learn robust representations for underrepresented subtypes and may inflate performance metrics for majority classes. Future work could explore data augmentation strategies or oversampling approaches tailored to rare leukemia subtypes. Moreover, the gradient-based interaction scores reported here reflect co-sensitivity of model outputs and are not direct measures of biological causality. Functional validation of the identified RNA-genomic feature pairs, for example through regulatory network analysis or experimental perturbation, will be necessary to translate these computational findings into mechanistic insights. Finally, the current framework integrates two omics modalities. Extension to three or more modalities (e.g., incorporating epigenomic or proteomic data) represents a natural and important future direction, and the GW-OT alignment module is in principle extensible to such settings.

## Conclusions

In summary, we proposed OTTER, a novel multi-modal deep learning framework that integrates transcriptomic and genomic data for T-ALL subtyping. By combining modality-specific encoders with GW-OT alignment, OTTER capture distinct statistical characteristics of gene expression and genomic variant data while preserving complementary relationships between the two omics. The improved classification performance, more discriminative fused latent representation, and results of the ablation analyses support the contribution of geometry-aware cross-omics alignment and variational regularization.

Gradient-based interpretation analysis further enables the identification of subtype-specific drivers and associated molecular programs, which are consistent with established T-ALL biology. The preservation of predicted subtype-associated structure in two independent clinical cohorts provides evidence for the generalizability of the learned representation. Overall, OTTER provides a computationally principled, biologically interpretable, and broadly applicable framework for multi-omics-driven cancer subtype classification, with direct implications for precision oncology and risk-stratified treatment of T-ALL. To facilitate reproducibility and broader application, we implemented our framework as an open-source python package, OTTER, for multi-omics integration and T-ALL subtype classification. The package is publicly available at https://github.com/wan-mlab/OTTER.

## Declarations

### Ethics approval and consent to participate

Not applicable

### Consent for publication

Not applicable

### Availability of data and materials

The datasets generated and/or analysed during the current study are available in the Synapse platform (ID: syn54032669; https://doi.org/10.7303/syn54032669). The NOPHO ALL2008 dataset is publicly available through the Gene Expression Omnibus under accession GSE272023, and the AIEOP dataset is provided from clinTALL in Supplementary Table S11.

## Authors’ contributions

L.L.: data preprocessing, machine learning model development, data analysis and interpretation, manuscript preparation, editing, and review. J.W.: manuscript editing and review. S.W.: study concept and design, manuscript editing and review.

## Acknowledgements

We thank the Children’s Oncology Group and the investigators involved in the AALL0434 study for generating and making available the clinical, transcriptomic, and genomic data analyzed in this work. We also acknowledge the investigators who generated the NOPHO ALL2008 and AIEOP-BFM ALL 2017 cohorts.

## Clinical trial number

not applicable

